# METI: Deep profiling of tumor ecosystems by integrating cell morphology and spatial transcriptomics

**DOI:** 10.1101/2023.10.06.561287

**Authors:** Jiahui Jiang, Yunhe Liu, Jiangjiang Qin, Jingjing Wu, Jianfeng Chen, Melissa P. Pizzi, Rossana L. Segura, Kohei Yamashita, Zhiyuan Xu, Guangsheng Pei, Kyung Serk Cho, Yanshuo Chu, Ansam F. Sinjab, Fuduan Peng, Guangchun Han, Ruiping Wang, Xinmiao Yan, Enyu Dai, Yibo Dai, Mingyao Li, Andrew Futreal, Anirban Maitra, Alexander Lazar, Xiangdong Cheng, Humam Kadara, Jaffer Ajani, Amir A. Jazaeri, Jianjun Gao, Jian Hu, Linghua Wang

## Abstract

The recent advance of spatial transcriptomics (ST) technique provides valuable insights into the organization and interactions of cells within the tumor microenvironment (TME). While various analytical tools have been developed for tasks such as spatial clustering, spatially variable gene identification, and cell type deconvolution, most of them are general methods lacking consideration of histological features in spatial data analysis. This limitation results in reduced performance and interpretability of their results when studying the TME. Here, we present a computational framework named, <u>M</u>orphology-<u>E</u>nhanced Spatial <u>T</u>ranscriptome Analysis Integrator (METI) to address this gap. METI is an end-to-end framework capable of spatial mapping of both cancer cells and various TME cell components, robust stratification of cell type and transcriptional states, and cell co-localization analysis. By integrating both spatial transcriptomics, cell morphology and curated gene signatures, METI enhances our understanding of the molecular landscape and cellular interactions within the tissue, facilitating detailed investigations of the TME and its functional implications. The performance of METI has been evaluated on ST data generated from various tumor tissues, including gastric, lung, and bladder cancers, as well as premalignant tissues. Across all these tissues and conditions, METI has demonstrated robust performance with consistency.

## INTRODUCTION

Spatial transcriptomics (ST) measures gene expression while persevering the spatial information that is not available in conventional single-cell RNA sequencing (scRNA-seq) [1]. The spatial location shed lights on TME’s cellular composition and organization, facilitating investigations into spatial gene expression patterns and cellular interactions at different tumor regions [2-4]. Commonly used ST platforms can be broadly categorized into two types: next-generation sequencing (NGS)-based, such as 10x Visium, GeoMx [5], Slide-Seq [6], and hybridization-based approaches such as MERFISH [7], seqFISH [8], and CosMx [9]. NGS-based ST methods cover the entire transcriptome but not at single-cell resolution, while in situ hybridization-based methods offer superior spatial resolution but are limited to a small portion of the genome, restricting their potential in discovery-based studies.

Many ST platforms allow a high-resolution scan of the Hematoxylin and Eosin stained (H&E) image of the same tissue section, which is valuable in downstream analysis. Various cell types, distinguished by their unique cell morphology, can be identified through close examination of the H&E image without the need for cell deconvolution analysis using gene expression. Also, potential technical and analytical artifacts can also be addressed more effectively by incorporating histological features into ST data analysis. Some state-of-the-art methods are developed to integrate spatial gene expression and images for various tasks. For example, MUSE [10] characterizes tissue composition through combined analysis of morphologies and transcriptional states for ST data using a deep learning approach; SpaGCN [11] utilizes a graph convolutional network (GCN) to integrate gene expression and histology to identify spatial domains; TESLA [12] uses convolutional neural network (CNN) to integrate gene expression and histology to map tumor core, edge, and different cell types at image pixel level. Despite the outstanding performance of these method, they share several limitations. Firstly, they are general analytic tools which works on data generated from any tissue type and are not specifically tailored for studying cancer cells and the TME. Without the ability to incorporate domain knowledge for cancer genomics, these methods may overlook crucial features specific to cancer cells and other key components in TME. Secondly, their reliance on “black box” machine learning models limits the interpretability of the results, as the models are solely data-driven without clear insights into underlying biological mechanisms. Lastly, these methods often neglect the investigation of associations between RNA expression and histology, which can hinder the discovery of important correlations between gene expression patterns and tissue morphology. These abovementioned limitations restrict our characterization of the key components and intricate interactions in TME.

In this study, we present a novel analytic framework that systematically analyzes cancer cells and cells of the TME by incorporating spatial gene expression, tissue histology, and prior knowledge of cancer and TME cells. Our methodology starts with the identification of key cellular components and their states within the TME, including various immune cells and their transcriptional states, tumor stromal components such as cancer-associated fibroblasts (CAFs), and the epithelial compartment. Meanwhile, METI offers complementary information on cell morphology for various cell type from the H&E images. The combined results from gene expression and histology features provide a comprehensive understanding of the spatial cellular composition and organization within the tissue. The evaluation of METI shows robust and consistent performance across ST datasets generated from diverse cancer types including gastric cancer, lung cancer, and bladder cancer (**Supplementary Table 1**).

## Results

### 1. Overview of METI’s workflow

METI analyzes the TME in a systematic, step-by-step manner, focusing on the progression from normal to premalignant cells, then to malignant cells, while also examining the lymphocytes within each tissue section. To achieve that, METI employs five parallel modules, each of which addresses a different component of the TME (**Fig. 1**). In the first module, METI identifies normal and premalignant cells, such as goblet cells in the stomach [13-15]. In module 2, METI identifies tumor cell-enriched regions and characterizes their cell states heterogeneity. Module 3 focuses on spatial mapping of T cells including CD4 and CD8 T cells, and various T cell states such as regulator T cells (Treg) and exhausted T cells (Tex). In addition to T cells, METI identifies other immune cells including neutrophils, B cells, plasma cells, and macrophages in module 4. In the last module, METI focuses on a comprehensive analysis of cancer-associated fibroblasts (CAFs), a subset of activated stromal cells that play a crucial role in cancer progression and therapy resistance [16-18]. This module maps CAFs and their subtypes, including myCAFs, iCAFs and apCAFs [19-23]. After identifying various cell type and cellular states in each module, METI next examines the relative spatial relationships of these components with cancer cells and investigate how the spatial coherence between different component affects their gene expression profiles.

**Fig. 1.**
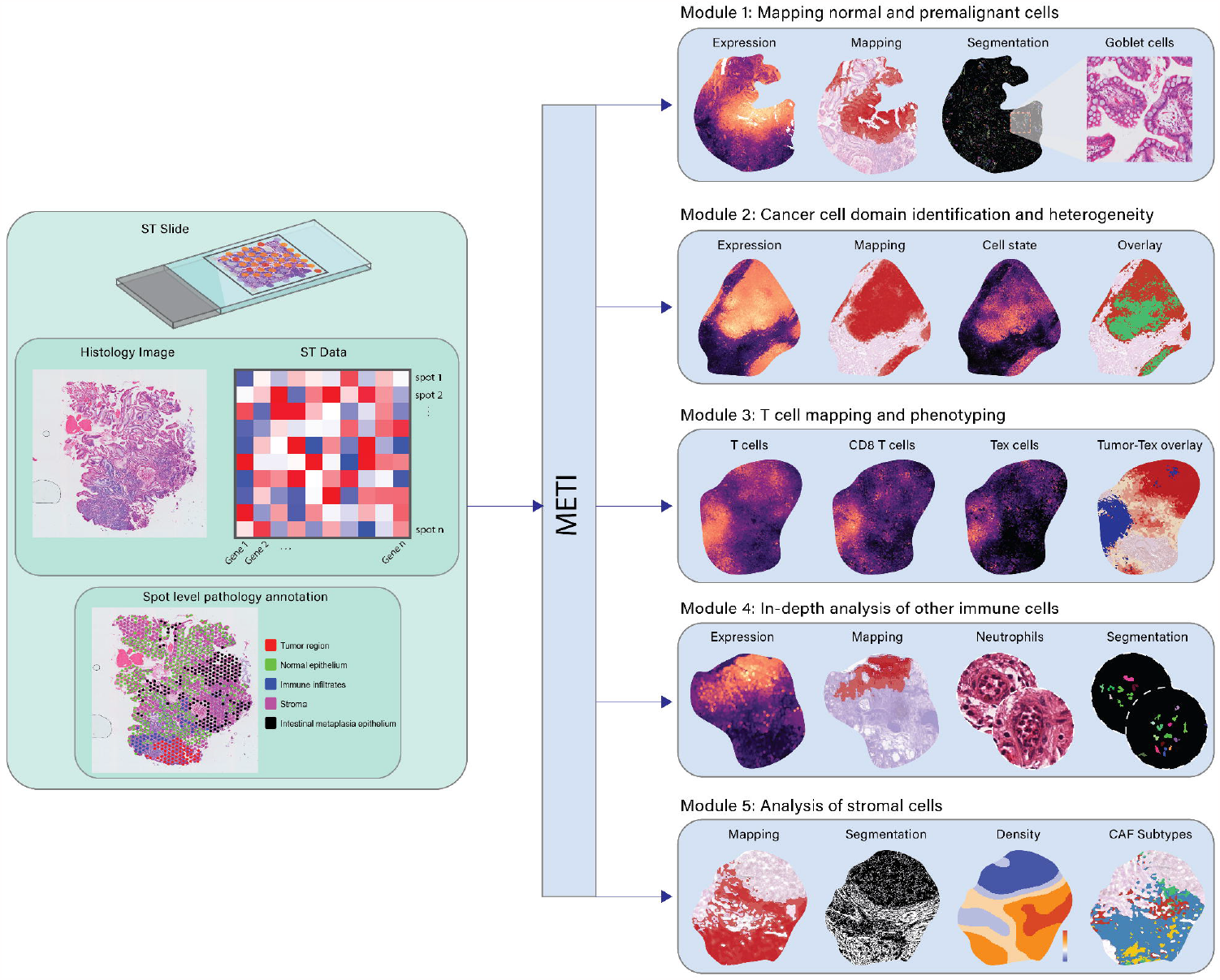
Workflow of METI. METI utilizes 10x Visium Spatial Transcriptomics (ST) data and original Hematoxylin and Eosin (H&E) images, in conjunction with pathology annotations. The framework encompasses five distinct modules. Module 1 is dedicated to mapping normal and premalignant cells through the integration of gene expression (GE) data and H&E images. Module 2 focuses on identifying cancer cell domains and characterizing their heterogeneity. Module 3 is dedicated to T cell mapping and phenotyping. Module 4 involves in-depth analysis of other lymphoid cells. Lastly, Module 5 pertains to the analysis of Cancer-Associated Fibroblasts (CAFs). Alongside cell type identification, METI offers nuclei segmentation and the functionality of generating 3D cell density plots.

### 2. Mapping normal and premalignant cells

The first module of METI focuses on dissecting the normal and premalignant cells within the epithelial cell compartment. We here used goblet cells as an example as they display a distinctive morphological appearance in H&E-stained images. Goblet cells are shaped like wine goblets, with pale, almost white vesicles at the top and oval nuclei at the base. Goblet cells are commonly found in the respiratory, digestive, and reproductive tracts, including the small intestine, colon, and bronchi. They play a critical role in maintaining homeostasis in these tissues. In the context of disease, the abnormal presence of goblet cells in the gut is a key characteristic of a precancerous condition of gastric cancer, known as intestinal metaplasia [13, 15, 24-26].

To demonstrate the power of METI’s module 1, we applied METI to identify goblet cells on a human stomach adenocarcinoma (STAD) sample labeled as G1. METI first combines canonical goblet cell markers reported in previous publications [27-31] into a meta gene signature, including *MS4A10, MGAM, CYP4F2, XPNPEP2, SLC5A9, SLC13A2, SLC28A1, MEP1A, ABCG2, and ACE2* (**Supplementary Table 2**). This meta gene visually represents the overall expression levels of the goblet cell molecular signature across the whole section. As shown in **Fig. 2b**, this meta gene was then used to annotate goblet cell-enriched regions using a machine learning model, TESLA [12]. Notably, cell type annotation in TESLA mainly relies on gene expression, which may lead to false negative annotation due to the regional variation and high level of noise for some marker genes. For instance, as shown in **Fig. 2c and 2e**, TESLA fails to identify goblet cells in region 4, which contains multiple goblet cells labeled by pathologists.Further examination shows that this false negative in detection was due to the overall low unique molecular identifier (UMI) counts captured in region 4, as illustrated in **Fig. 2d**. To address the limitations caused by low-quality gene expression data, METI simultaneously performed goblet cell identification on the H&E image. METI employed a K-mean-based segmentation method to detect different morphological components, such as background, nuclei, fiber, gland, and necrosis. Next, by filtering on the color, shape, and size of these components (see Methods), METI is able to accurately detect individual goblet cells characterized by their morphology signature, i.e., their round hollow centers (**Fig. 2e)**. Then, METI combined the goblet cells identified from gene expression and image as the final results. This integrative approach for goblet cell detection overcomes the limitations posed by low UMI counts and provides a more accurate characterization of goblet cells within the analyzed samples. Although we showcase the capability of this module in goblet cell detection, this framework can easily be adapted to identify other normal and premalignant cells using both transcriptomics and histology images.

**Fig. 2.**
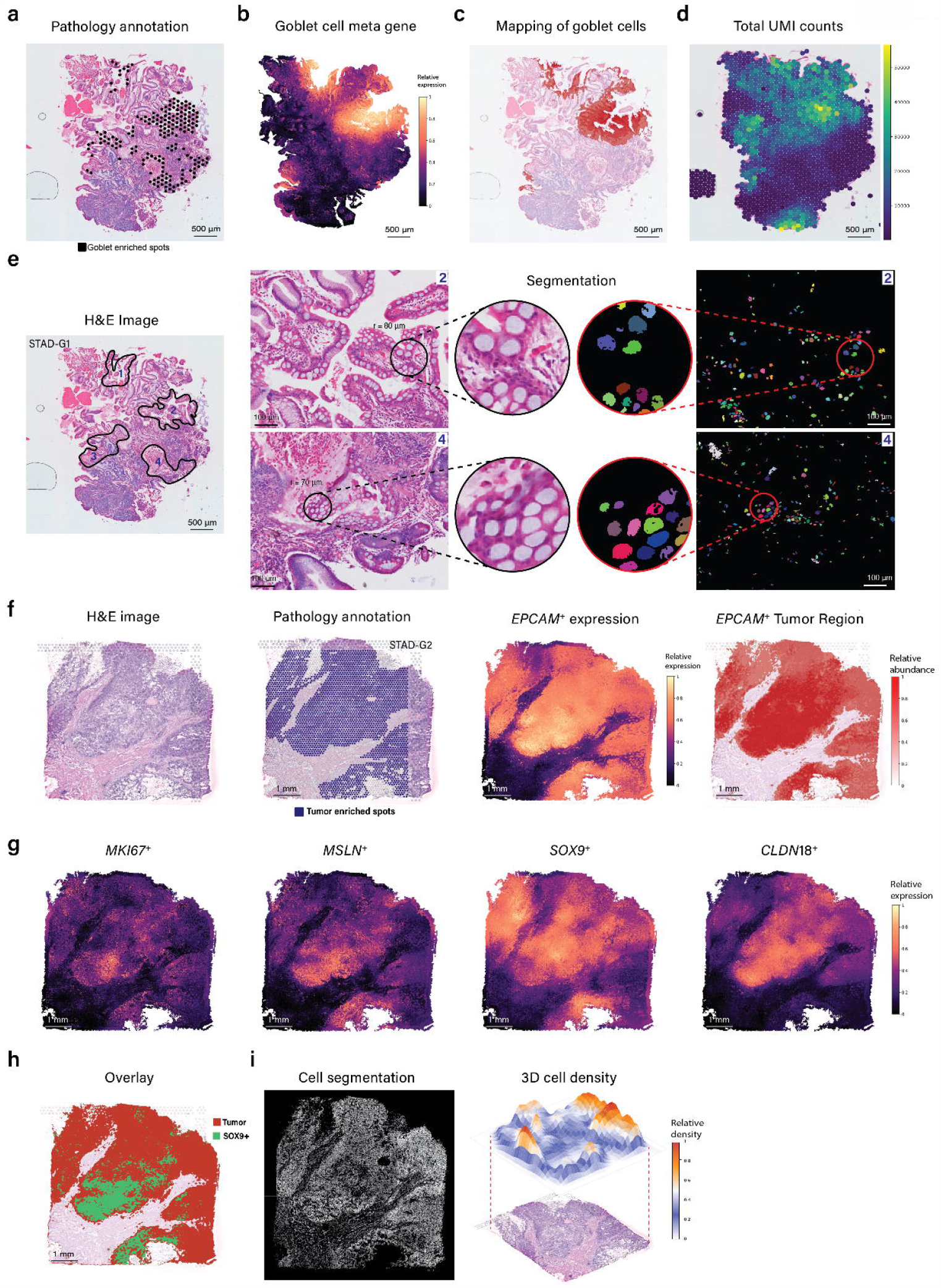
Mapping premalignant cells and cancer cell domain. **a**, Pathology annotation depicting goblet cell enriched spots in STAD G1. **b**, Goblet meta gene expression plot at pixel-level. **c**, Annotation indicating regions of high goblet cell gene expression on the H&E image. **d**, Total UMI counts for individual spots. **e**, Identification of four distinct goblet-enriched regions on the left side, accompanied by zoom-in views of goblet regions of H&E image and segmentation outcomes for regions 2 and 4. **f**, Pathology annotation highlighting tumor cell enriched spots of STAD G2 (left), pixel depiction of *EPCAM* highly expressed regions (middle), and *EPCAM*^*+*^ region annotation on the H&E image (right). **g**, Pixel-level gene expression plots for tumor subtypes, *MKI67, MSLN, SOX9*, and *CLDN18*. **h**, Overlay of regions expressing tumor-related genes and *SOX9*-positive regions. **i**, Nuclei segmentation (left) alongside 3D cell density plots (right).

### 3. Identification of Cancer Cell Domains and Heterogeneity

The majority of solid tumors originate from epithelial cells, known as carcinomas, including gastric, lung, bladder, breast, prostate, and colon cancers, while some other solid tumors start in other types of tissues including sarcoma and melanoma. Regardless of their cells of origin, understanding the molecular features and cellular heterogeneity of malignant cells is crucial for unraveling the mechanisms underlying tumor growth, invasion, metastasis, and therapeutic response. Therefore, METI’s second module focuses on the analysis of malignant cells. This module starts by identifying cancer cells using cancer cell markers that are curated by the authors such as cytokeratins (CK), *EPCAM*, and trefoil factors (TFF). As depicted in **Fig. 2f**, METI effectively identifies all tumor regions in STAD sample G2, in strong agreement with annotations made by our experienced pathologists. Next, METI incorporates additional markers to characterize cancer cell states and heterogeneity, including markers of cell proliferation such as *MKI67* to map proliferative cancer cells, stemness-related markers such as *SOX9* to identify stem-like cancer cells in STAD, and therapeutic targets like *CLDN18* and *MSLN* to further characterize tumor subtypes [32-35]. These aforementioned marker genes exhibit distinct expression patterns within the tumor region of sample G2, as illustrated in **Fig. 2g**. They can be utilized to characterize different states of cancer cell states. For example, as shown in **Fig. 2h**, METI is not only able to identify the *SOX9*^*+*^ tumor region, but also can illustrate the co-localization or exclusivity of different cancer cell states in **Supplementary Fig. 1**. This module offers a flexible and customizable approach, allowing users to input their genes of interest for tumor state identification. Additionally, users can employ genes associated with critical pathways such as *KRAS, EGFR*, and factors like hypoxia to conduct a comprehensive exploration of cancer cell states and spatial heterogeneity across diverse cancer types.

Quantifying the distribution and density of cells spatially within biological tissues is crucial for diverse applications, particularly in the field of pathology and oncology. While gene expression provides a molecular lens, the associated H&E images can be leveraged to measure spatial cell distribution and density. Following a parallel process in module 1, METI next conducted tumor cell nuclei segmentation, and then generated 3D tumor cell density plots (**Fig. 2i**), visually depicting the spatial distribution and density of cancer cells. This function serves to convey the spatial distribution, density and pattern of cell types of interest.

### 4. T Cell mapping and phenotyping

Module 3 in METI is dedicated to characterizing T cells and their various states within the TME. Initially, we utilize specific T cell markers, including *CD3D* and *CD3E*, to map T cell-enriched regions. Within the identified T cell regions, we further discern the different states of T cells. By adding specific cell lineage markers such as *CD4, CD8A*, and *CD8B* [36], we can further distinguish CD4 T cells, CD8 T cells, and their various states including CD4^+^ Tregs (e.g., *FOXP3, IL2RA*) and CD8^+^ Tex cells by incorporating known immune checkpoint genes (e.g., *PD-1, TIM-3*, and *LAG-3, CTLA-4, TIGIT*) and Tex related transcription factors (e.g., TOX) [36]. Furthermore, this module provides function of overlaying two or more different T cell states within defined cancer cell regions directly on the same tissue section, allowing us to visualize their spatial relationships. Given that the level and spatial distribution of infiltrated T cells are both critical factors influencing tumor immune phenotypes and immunotherapy responses, METI’s 3D module creates cell density plots for the entire image, serving to visually depict the spatial distribution of T cells within the TME.

To showcase the capability of this module, we applied METI to analyze a STAD sample G3 and a lung adenocarcinoma (LUAD) sample L1. METI discerned regions characterized by elevated T cell gene expression levels, as illustrated in **Fig. 3a**. Next, to delineate regions enriched in CD4 T cells and CD8 T cells, we restrict our analysis to T cell-enriched regions only. This approach was undertaken to specifically identify spots with CD4 T cells and differentiate them from macrophages, both of which can express *CD4*. The regions enriched in CD4 T cells and CD8 T cells are shown in **Fig. 3b**, and their different states including CD4^+^ Tregs and CD8^+^ Tex cells are shown in **Fig. 3c and 3d**. Mapping distinct T cell states aids in elucidating their spatial landscape and relationships within the analyzed STAD and LUAD samples, as well as cellular interactions, fostering the generation of insightful hypotheses.

**Fig. 3.**
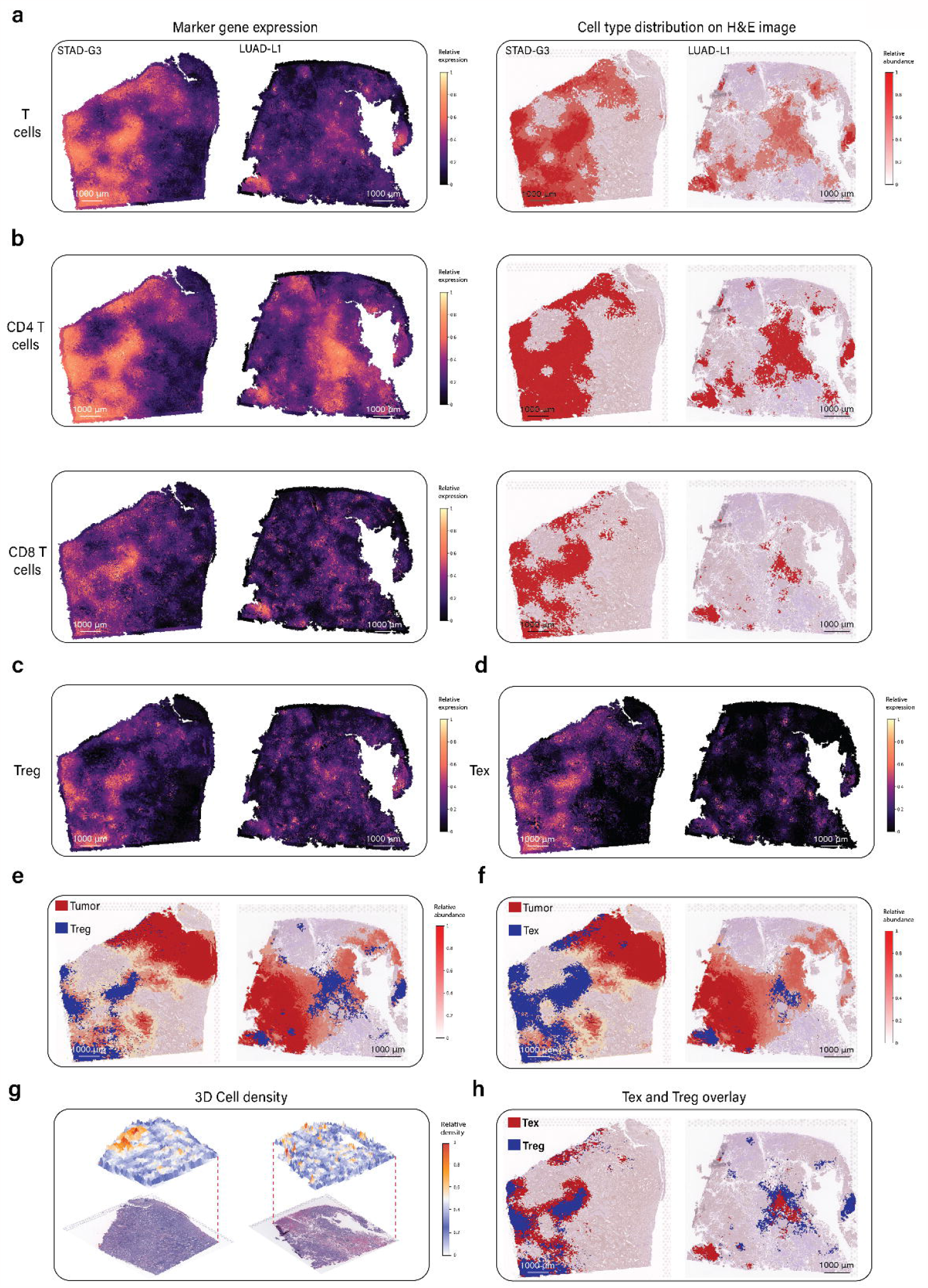
T Cell mapping and phenotyping. **a**, Pixel-level visualization of T cell marker gene expression in STAD G3 and LUAD G1 (left), accompanied by annotation indicating regions of T cell marker gene expression on the H&E image. **b**, Pixel-level representation of CD4 T cell and CD8 T cell marker gene expression (left), along with annotation of CD4 T cell and CD8 T cell marker gene-expressing regions on the H&E image (right). **c**, Pixel-level presentation of CD4+ Treg marker gene expression. **d**, Pixel-level depiction of CD8^+^ Tex marker gene expression. **e**, Overlay displaying the intersection of tumor+ region and CD4^+^ Treg positive region. **f**, Overlay illustrating the overlap between tumor+ region and CD8^+^ Tex positive region. **g**, 3D cell density plots for STAD G3 and LUAD G1. **h**, Overlay demonstrating the spatial relationship between CD4^+^ Treg and CD8^+^ Tex positive regions.

As the relative locations of CD4^+^ Tregs and CD8^+^ Tex cells to cancer cells impact tumor immune phenotypes [37, 38] and immunotherapy responses, we have overlaid regions with cancer cells with those enriched with CD4^+^ Tregs and CD8^+^ Tex cells, respectively, as depicted in **Fig. 3e and 3f**. Based on the overlay results, we observe distinct enrichment patterns in CD4^+^ Tregs and CD8^+^ Tex cells across different samples. Specifically, in sample G3, CD4^+^ Tregs are slightly less abundant than CD8^+^ Tex cells. Conversely, in the sample L1, CD8^+^ Tex cells are less abundant than CD4^+^ Tregs (**Fig. 3e-f**). This highlights the variability in T cell states within different tumor types. To better illustrate the spatial cell distribution of the whole image, METI provides 3D cell density plots (**Fig. 3g**) based on the nuclei density segmented from the H&E image. For the STAD sample, a region in the upper left displays higher cell density, whereas the LUAD sample shows relatively homogeneous cell density throughout. Furthermore, we conducted an overlay of CD4^+^ Treg and CD8^+^ Tex signals to study their spatial co-localization patterns (**Fig. 3h**). Notably, CD4^+^ Tregs and CD8^+^ Tex cells tend to co-localize at the bottom left of the LUAD sample, while the rightmost part of the LUAD sample solely comprises CD4^+^ Treg cells, indicating the heterogeneity in spatial distribution and cellular composition of T cells. This co-localization analysis provides a better understanding of the coexistence and potential interplay between these two T cell states. Moreover, METI can assist researchers in studying various types of T cells, such as naïve T cells, memory T cells, follicular helper T cells and their transcriptional states. Users can customize it to plot specific T cell types and states of interest. This flexibility allows researchers to explore T cell state composition and distribution within TME.

### 5. In-depth analysis of other immune cells

METI’s module 4 is capable of detecting immune cell types other than T cells that are critical components in the TME including neutrophils, macrophages, B cells, and plasma cells. METI utilizes validated gene signatures to identify specific immune cell types/states [36, 39-41]. We have applied this module to two bladder cancer samples B1 and B2 for neutrophil detection. The two H&E images are shown in **Fig. 4a and 4c**. The neutrophil-enriched regions in these two sections were verified by our experienced pathologists as ground truth for evaluation (**Supplementary Fig. 3**). As shown in **Fig. 4b and 4d**, METI identified regions exhibiting elevated gene expression levels of neutrophil marker genes in both sections. Subsequently, METI conducted corresponding annotation for neutrophil-enriched regions directly on H&E image, isolating regions expressing the neutrophil marker genes and providing a magnified view, as illustrated in **Fig. 4e**. Upon zooming in on these annotated regions, neutrophils, which exhibit characteristic multi-lobed nuclei, were easily distinguished in the image analysis. In **Fig. 4f**, four regions where neutrophils have been pathology-verified were circled out and subsequently segmented for neutrophil detection. The results correlated well with the annotation using gene expression (**Supplementary Fig. 3**). We also provided 3D cell density as shown in **Fig. 4g** to illustrate the spatial cell distribution in the vicinity of neutrophils within the TME.

**Fig. 4.**
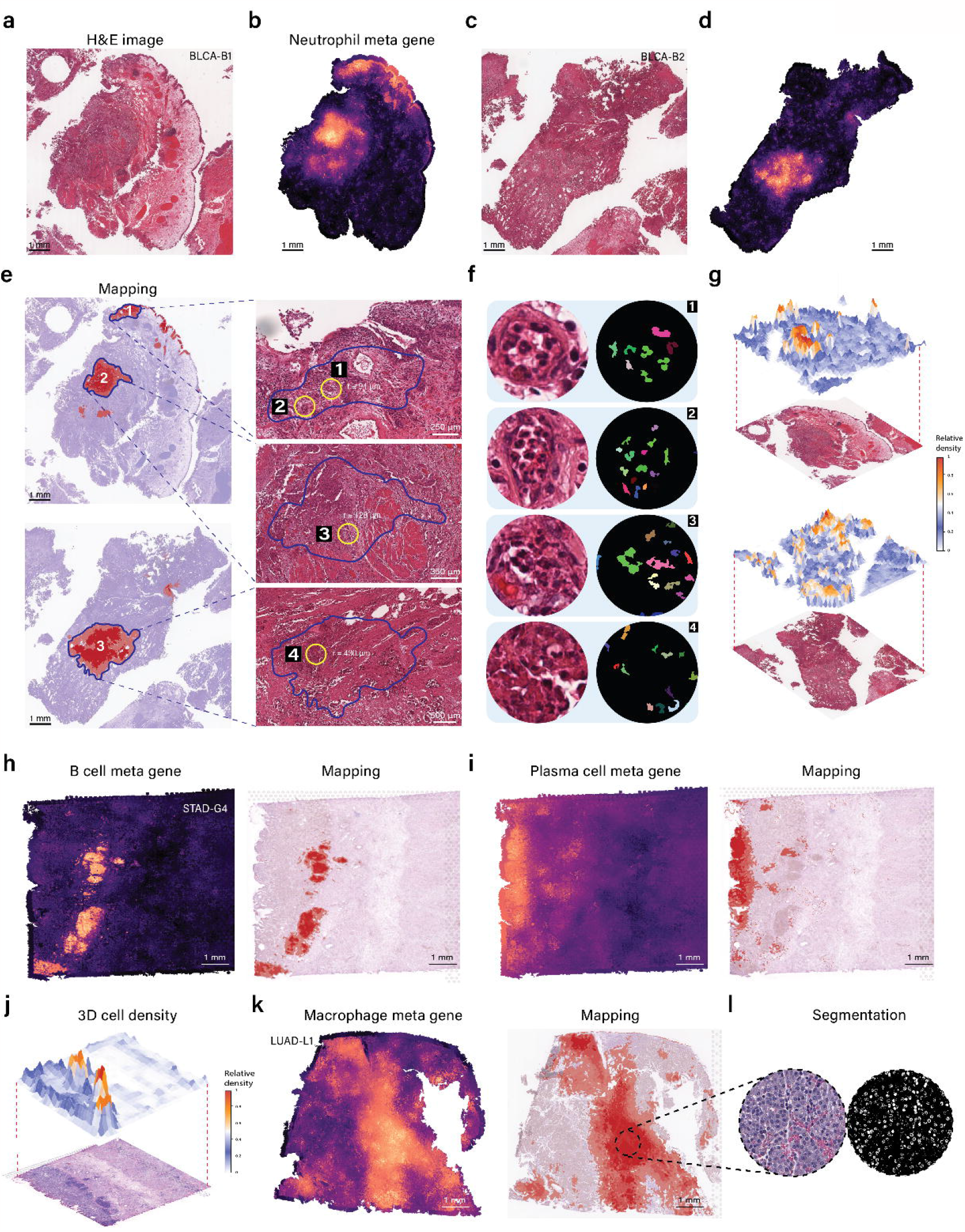
In-depth analysis of other lymphoid cells. **a**, H&E image of bladder cancer sample B1. **b**, Pixel-level visualization of neutrophil marker gene expression in BLCA-B1. **c**, H&E image of bladder cancer sample BLCA-B2. **d**, Pixel-level visualization of neutrophil marker gene expression in BLCA-B2. **e**, Annotation indicating regions of high neutrophil gene expression on the H&E image for BLCA-B1 and BLCA-B2; Zoom-in display of three neutrophil enriched region of BLCA-B1 and BLCA-B2, and four yellow circled regions where neutrophils present visually. **f**, zoom-in view of four yellow circled region in **e** and corresponding segmentation result. **g**, 3D cell density plots for BLCA-B1 and BLCA-B2. **h**, Pixel-level visualization of B cell marker gene expression in STAD G4 (left), accompanied by annotation indicating regions of B cell marker gene expression on the H&E image (right). **i**, Pixel-level visualization of plasma cell marker gene expression in STAD G4 (left), accompanied by annotation indicating regions of plasma cell marker gene expression on the H&E image (right). **j**, 3D cell density plots for STAD G4. **k**, Pixel-level visualization of macrophage marker gene expression in STAD G4 (left), accompanied by annotation indicating regions of plasma macrophage marker gene expression on the H&E image (right). **l**, Zoom-in view of macrophage regions of H&E image and segmentation.

Additionally, we demonstrate the capability of this module by mapping B cells and plasma cells on an STAD sample (**Fig. 4h and 4i**). The 3D cell density plot (**Fig. 4j**) aligns well with the lymphoid aggregates in sample STAD G4 annotated by our pathologists (**Supplementary Fig. 2a**). Similarly, macrophages can be correctly mapped in the LUAD sample based on pathology annotation (**Fig. 4k and Supplementary Fig. 2b**). Within the regions showing high macrophage marker gene expression, a randomly selected region is segmented, revealing a cluster of macrophages (**Fig. 4l**). In addition to the aforementioned immune cell types, this module maintains flexibility by allowing users to investigate other specific immune cell populations of interest using their curated gene signatures.

### 6. Analysis of Cancer-Associated Fibroblasts (CAFs)

In Module 5, METI is designed to analyze stromal cell components including CAFs and various CAFs subtypes within the TME. CAFs are known for their exceptional heterogeneity, both phenotypically and functionally [42-44]. They are categorized as activated fibroblasts, representing an essential component of the TME with both tumor-promoting and tumor-restraining activities [45-47]. CAFs are phenotypically and functionally heterogeneous. Different subtypes of CAFs such as myofibroblastic CAFs (myCAFs), inflammatory CAFs (iCAFs), and antigen-presenting CAFs (apCAFs) have been identified and described [19-23].

We applied module 5 to the gastric sample G2 which was annotated to contain abundant tumor stroma by our pathologists (**Fig. 5a**). METI first segmented CAFs and generated a fibroblast cell density plot as illustrated in **Fig. 5b**. Next, we found that the fibroblasts enriched region annotated by METI have diminished UMI counts, aligning with the notion that cancer cells tend to have higher UMI counts compared to other cell types (**Fig. 5c**). METI next effectively mapped CAFs within the sample using the CAF metagene (**Fig. 5d**) and annotated CAFs directly on H&E image, which was highly consistent with the pathology annotation. Within the annotated CAF regions, METI further delves into the characterization of CAF subtypes, including myCAFs, iCAFs, and apCAFs (**Fig. 5e-g**) [19-23]. To characterize the spatial colocalization of CAF subtypes, we overlayed the three CAF populations with the total CAF-positive regions (**Fig. 5h**).This approach allows us to better understand the spatial heterogeneity of CAFs within the TME. Likewise, METI can co-map CAFs, cancer cells, and any other immune cell subsets of interest to provide additional insights into cellular interactions among them. This module remains adaptable, enabling users to explore other subregions related to CAFs based on their specific interests.

**Fig. 5.**
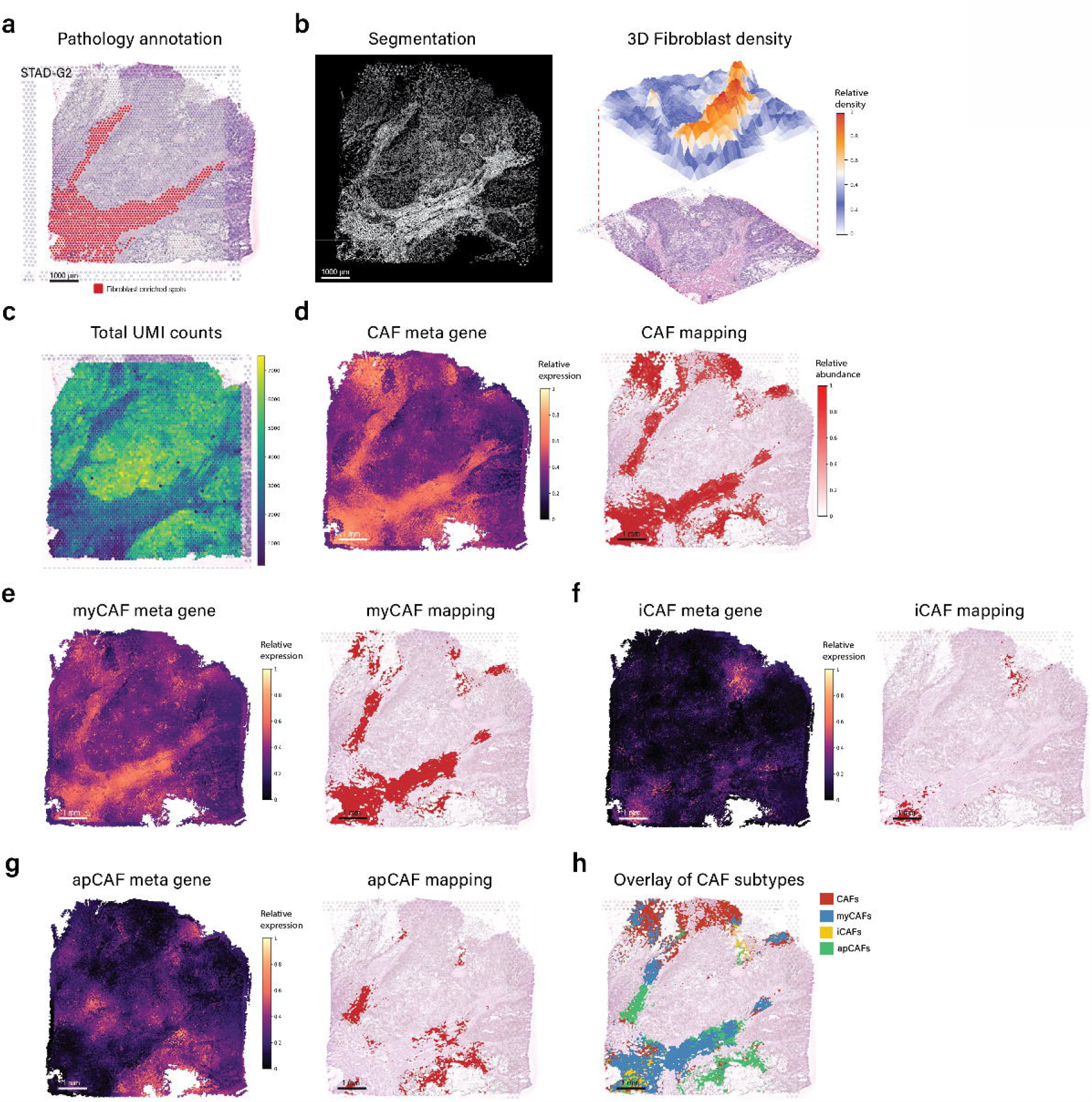
Analysis of Cancer Associated Fibroblasts. **a**, Pathology annotation of fibroblast enriched spots in STAD G2. **b**, Fibroblasts segmentation result (left), accompanied by 3D fibroblast density plots (right). **c**, Total UMI counts for individual spots. **d**, Pixel-level meta gene expression plot for CAF (left), with annotations highlighting regions of elevated CAF gene expression on the corresponding H&E image (right). **e**, Pixel-level meta gene expression plot specifically for myCAF (left), accompanied by annotations indicating regions of elevated myCAF gene expression on the H&E image (right). **f**, Pixel-level meta gene expression plot for iCAF (left), with annotations denoting regions of elevated iCAF gene expression on the H&E image (right). **g**, Pixel-level meta gene expression plot for apCAF (left), with annotations indicating regions of elevated apCAF gene expression on the H&E image (right). **h**, Overlay showcasing regions of high gene expression for myCAF, iCAF, apCAF, and general CAF.

## Discussion

In this study, we present METI, a robust machine-learning framework designed to meet the need for comprehensive profiling of various cell types and cellular states within the TME. By effectively integrating transcriptomics and histological image information, METI decreases the risk of non-specific mapping of various key cell types in TME. This ability sets METI apart from current cell deconvolution tools that primarily rely on gene expression rather than histological morphology.

METI comprises five modules that characterize various cellular components, encompassing distinct types and states of tumor cells, immune cells, and stromal cells. By analyzing transcriptomics and histological information jointly, METI provides a more comprehensive approach to understanding cancer cells and the TME compared to analyzing the gene expression and histology images individually. Our results have shown that METI can effectively identify cells present in the tissue regions with low-quality gene expression. This integrative approach enables researchers to maximize the information extracted from their data. In addition to general pattern extraction, METI is uniquely capable of stratifying various cell states, including CD4^+^ Tregs, CD8^+^ Tex cells, iCAFs, myCAFs, apCAFs, among many others. Notably, this stratification is not restricted to the aforementioned subtypes; users have the flexibility to define and explore other cell subtypes to suit their research needs. METI has been evaluated using different cancer datasets (**Supplementary Table 1**), including samples from lung adenocarcinoma, bladder cancer, and gastric cancer for which we have obtained spot-level pathology annotations as the ground truth for the assessment of its performance. METI stands out by characterizing both known and custom-defined cell populations. This adaptability and potential for tailored exploration make it a powerful tool for personalized research.

While METI presents a useful framework for in-depth cancer cell and TME cell profiling, it is important to recognize its limitations. One notable challenge arises from the dependence on transcriptomics data, which can be susceptible to technical fluctuations, constrained sample quality, dropout occurrences, and dependence on user-provided gene signatures, possibly resulting in the generation of false negative outcomes. Moreover, the performance of METI’s segmentation function could exhibit variability contingent upon the quality and resolution of the images being analyzed.

## Acknowledgement

This study was supported in part by the National Institutes of Health/NCI grants R01CA266280 (to L.W.), U01CA264583 (to H.K. and L.W.), the start-up research fund, and the University Cancer Foundation via the Institutional Research Grant Program at the University of Texas MDACC and The Break Through Cancer Foundation (to L.W., A.A.J., and A.M.), as well as the Cancer Prevention and Research Institute of Texas award RP220101 (to H.K. and L.W.). In addition, A.M. was also supported by the Sheikh Khlaifa bin Zayed Foundation and the MDACC Moon Shots Program in Pancreas Cancer. L.W. and H.K. are Andrew Sabin Family Foundation Fellows at MDACC. A.S. is supported by an NCI T32CA217789 MDACC postdoctoral fellowship. This study was also supported by MDACC Support Grant (CA016672).

## Author contribution

L.W. conceived the study and designed the experiments. L.W. and J.H. jointly supervised the study. J.J.Q., J.W., J.C., M.P.P., X.C., A.S., Z.X., H.K., K.Y., J.A., and J.G. contributed to sample and information collection and data generation. R.L.S., J.W., A.L., reviewed the histology images and performed pathology annotation. L.W. supervised the bioinformatics data processing and analysis. J.J. performed bioinformatics data analysis and developed METI. Y.L., Y.C., F.P., G.H., R.W., X.Y., E.D., and Y.D. assisted with data processing and analysis. L.W., J.H., J.J., M.L., H.K., J.A., A.A.J., and J.G. contributed to data interpretation. J.J., J.H., and L.W. wrote and revised the manuscript and all co-authors reviewed the manuscript.

## Conflict of interest statement

The authors declare no competing interests.

## Data and code availability

A detailed description of lung cancer and gastric Visium datasets including data sources and accession numbers RE described in our previous studies [36, 39]. The lung cancer (LC_1) dataset [48] can be downloaded from EGA under accession number EGAS00001005021. The gastric cancer and bladder cancer datasets will be uploaded and make publicly accessible upon the acceptance of the manuscript. All original code has been deposited at GitHub (https://github.com/Flashiness/METI) and is publicly available as of the date of publication.

## Methods

### Human samples

Two patients with primary muscle-invasive bladder carcinoma (T2–T4) were evaluated at the MD Anderson Cancer Center and underwent trans urethral resection of bladder tumor (TURBT). All patients received no treatments at the time of surgery. Formalin-fixed paraffin embedded (FFPE) samples in this study were obtained from Pathology Department under a waiver of consent from banked tissues approved by MD Anderson institutional review board protocols.

### ST Data Generation

ST on FFPE slides were performed with the Visium spatial technology from 10X Genomics. Two to three consecutive tissue sections of 5-μm thickness were collected for RNA extraction with the Qiagen RNeasy FFPE Kit. To assess the RNA quality of the tissue, the purified RNA was immediately processed to calculate the percentage of total RNA fragments >200 nucleotides (DV200) using the Agilent RNA 6000 Pico Kit. Based on DV200 evaluation, blocks with DV200 >30% were selected for proceeding with sectioning. The area of interest (11 x 11 mm) on section was carefully placed within the allowable area to ensure compatibility with the Visium CytAssist instrument. The tissues were then deparaffinized, stained, and decross-linked, followed by probe hybridization, ligation, CytAssist enabled RNA digestion and oligo capture, release, and extension. The Visium spatial gene expression FFPE libraries were constructed using the Visium CytAssist Spatial Gene Expression for FFPE Human Transcriptome Probe Kit (PN-1000444) following the manufacturer’s guidance. Constructed libraries were sequenced on the Illumina NovaSeq 6000 platforms to achieve a depth of at least 75,000 mean read pairs per spot.

### Data Processing

METI takes spatial gene expression and histology image data as input. The ST gene expression data contains an *N × M* matrix of unique molecular identifier (UMI) counts, where *N* denotes the number of spots and *M* represents the number of genes. Each spot is associated with 2-dimensional spatial coordinates denoted as (x, y). The gene expression values for each spot are normalized by dividing the UMI count of each gene within that spot by the overall UMI count of all genes in the same spot. The result is then scaled up by a factor of 10,000 and converted to a natural logarithm scale.

### Meta Gene plot generation

We utilized TESLA to get a pixel-level gene expression matrix. For a cell type which has *K* marker genes, we combine these *K* markers into a single meta marker gene. Considering that not all marker genes may be expressed, the meta gene is designed to preserve the expression patterns for at least a subset of the marker genes. Given *K* marker genes and a predefined number *k* based on the specificity of the markers, *k ≤ K*, for any given pixel *i*, we first sort the relative expression of all markers in descending order as {*e*_*1,i*,_ *e*_*2,i*,_ *e*_*3,i*,….,_ *e*_*k,i*_} Next, we select the top *k* expression values and compute the meta gene’s relative expression at pixel *i* as:

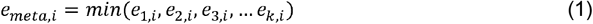

If number of expressed marker genes less than *k* which is *k*^th^ marker has zero expression at a given pixel, the meta gene’s relative expression will be 0 at that pixel. This ensures that expression patterns present in less than *k* genes are excluded from the meta gene, preventing the generation of less representative patterns.

### T cell states mapping

First, we employed *CD3D* and *CD3E* as markers to accurately annotate T cell-enriched regions using TESLA. The output of TESLA is a binary value array *A*_*T* cell_ of the same size as the histology image, in which value of 1 indicates T cell enrichment while 0 indicates non-enrichment of T cells. Similarly, we used *CD4* marker gene to identify rough CD4 T cell distribution patterns using TESLA, and the results is stored as a binary array 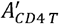 indicating the enrichment of rough CD4 T cells. Since CD4 T cells are a subset of T cells, we then filter 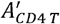 based on *A*_*T cell*_ to get a more accurate enrichment matrix of CD4 T cell, *A*_*CD*4*T*_, as below:

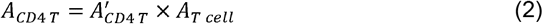

This operation filtered out false positive detected CD4 T cells by TESLA which do not express general T cell markers. Using a similarly pipeline, METI is able to identify CD8 T cells using *CD8A* and *CD8B*. We obtained array 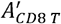 where 1 denotes *CD8A* or *CD8B* positive and 0 denotes negative. By intersecting *A*_*T cell*_ and 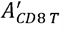, we obtained *A*_*CD*8*T*_ storing pixel with CD8 T cell positive expression.

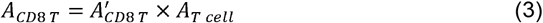

Next, to identify the CD4^+^ Treg cells, we input specific marker genes for Tregs, including *FOXP3* and *IL2RA*, to delineate the corresponding regions and generate an array 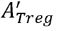. Given that CD4^+^ Treg cells is a subset of CD4 T cells, we construct a new binary array 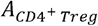, by intersecting *A*_*CD*4*T*_ and 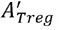. By visualizing *A*_*Treg*_, we obtained the annotation that outlines the CD4^+^ Treg signature on the H&E image.

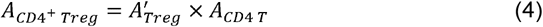

The identical procedure is employed to identify CD8^+^ Tex cells. We input specific marker genes for Tex cells, including *HAVCR2, LAG3, CTLA4, TIGIT, PDCD1*, and *LAYN*. With prior knowledge [49], any pixel expresses any 2 marker genes out of these 6 marker genes will be set to 1. To annotate the corresponding regions, we generate an array 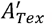. Like the CD4^+^ Treg identification process, we create a new binary array 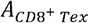 for CD8^+^ Tex cells by intersecting A*CD*8*T* and 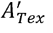. This array annotates the CD8^+^ Tex cell signature on the H& E image. In this case, we have *K* = 6 and set *k*= 2 based on previous publication, we derived the meta gene as formula (1), in which pixels with at least 2 marker genes expression out of the 6 have non-zeros expression. Then 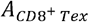 can be obtained as:

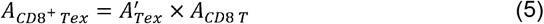

For other immune cell annotations, the markers and choice of *k* can be found in **Supplementary Table 3**.

### Nuclei segmentation

We first divided the image into multiple patches of size 2000 × 2000 pixels, and supplement incomplete patches on the border with black blocks to 2000 × 2000 pixels. We next converted the image from the default blue (B), green (G), red (R) color format to RGB format using OpenCV and then reshaped the 3D image array into a 2D array, where each row represents a pixel in the image, and three columns corresponds to the RGB color values of each pixel.Subsequently, we employed OpenCV K-means clustering algorithm to 2D image array with 10 channels, yielding two crucial outcomes, “centers” storing the coordinates of 20 centroids representing the color of each centroid, and “labels” storing the labels for each pixel indicating which centroid it belongs to. By assigning colors from the “centers” array to each pixel in the image based on their cluster labels stored in the “labels” array, segmented images were created. To better visualize the K-means clustering results, we set all the cluster centers to black as initial values and plot each color channel separately. To identify the cluster corresponding to the targe region, such as nuclei, we performed a color mapping. Users first need to input the color of target region in RGB format based on the prior knowledge, for example:

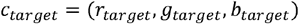

For each cluster derived from K-means segmentation, we calculate its average color value as:

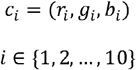

Next we calculate the Euclidean distance of each centroid color to *c*_*target*_ :

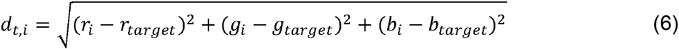

for each *i* ∈{1,2,…,10}. Then we look for the cluster with the minimum Euclidean distance:

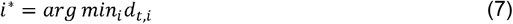

where *i*^*\**^ represents the index of the cluster that is closest in color to *c*_*target*_.

For example, in immune cell nuclei detection, we can use (50, 0, 100) as they appear dark blue/purple on H&E image, while for goblet, we used (240, 230, 230) as they look pale and clear. It is possible for a channel contains similar-colored noisy elements. We then incorporated a closing morphological operation using “morphologyEx” in OpenCV on the segmented results as a refinement step. This operation involves a combination of dilation and erosion processes as below:

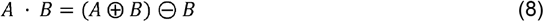

where ⨁ and ⊖ denote the dilation and erosion, respectively.

This step aims to eliminates noise points, while concurrently dilate the nucleus. Subsequently, the nucleus is returned to its normal size through the erosion process. For certain process such as filtering our goblet cells, we have one more step applying filters to refine the result which helps us to keep specific cells only. For instance, with filters such as an element’s area > 40, solidity > 0.5, and a length-to-width ratio < 3, we were able to distinguish goblet cells from other irrelevant elements. Notably, these parameter values are not fixed while they are adaptable based on cell types and samples. We employed a similar segmentation process for immune cells and fibroblasts, with the key distinction being the use of different filters and parameter values tailored to each cell type.

### 3D cell density visualization

Utilizing the nuclei detected from the previous step, we further performed a strong erosion to make the overlapped nuclei separable, follow by objective detection using function “findContours” in OpenCV. Then we computed the density of nuclei within a 20 × 20-pixel patch by using the spatial coordinates of each spot. These nuclei counts were served as the Z-coordinate values to construct our density plot. We generated a 3D surface map based on the complete set of Z-values.

## Notes

https://github.com/Flashiness/METI

### Competing Interest Statement

The authors have declared no competing interest.

## References: (please add relevant references in the main text)

1. Larsson, L., J. Frisen, and J. Lundeberg, Spatially resolved transcriptomics adds a new dimension to genomics. Nat Methods, 2021. 18(1): p. 15–18.

2. Walker, B.L., et al., Deciphering tissue structure and function using spatial transcriptomics. Commun Biol, 2022. 5(1): p. 220.

3. Baccin, C., et al., Combined single-cell and spatial transcriptomics reveal the molecular, cellular and spatial bone marrow niche organization. Nat Cell Biol, 2020. 22(1): p. 38–48.

4. Hwang, W.L., et al., Single-nucleus and spatial transcriptome profiling of pancreatic cancer identifies multicellular dynamics associated with neoadjuvant treatment. Nat Genet, 2022. 54(8): p. 1178–1191.

5. Merritt, C.R., et al., Multiplex digital spatial profiling of proteins and RNA in fixed tissue. Nat Biotechnol, 2020. 38(5): p. 586–599.

6. Rodriques, S.G., et al., Slide-seq: A scalable technology for measuring genome-wide expression at high spatial resolution. Science, 2019. 363(6434): p. 1463–1467.

7. Chen, K.H., et al., RNA imaging. Spatially resolved, highly multiplexed RNA profiling in single cells. Science, 2015. 348(6233): p. aaa6090.

8. Lubeck, E., et al., Single-cell in situ RNA profiling by sequential hybridization. Nat Methods, 2014. 11(4): p. 360–1.

9. Beechem, J.M., High-plex Multiomic Analysis in FFPE at Subcellular Level by Spatial Molecular Imaging.

10. Bao, F., et al., Integrative spatial analysis of cell morphologies and transcriptional states with MUSE. Nat Biotechnol, 2022. 40(8): p. 1200–1209.

11. Hu, J., et al., SpaGCN: Integrating gene expression, spatial location and histology to identify spatial domains and spatially variable genes by graph convolutional network. Nat Methods, 2021. 18(11): p. 1342–1351.

12. Hu, J., et al., Deciphering tumor ecosystems at super resolution from spatial transcriptomics with TESLA. Cell Systems, 2023. 14(5): p. 404-+.

13. Correa, P., M.B. Piazuelo, and K.T. Wilson, Pathology of gastric intestinal metaplasia: clinical implications. The American journal of gastroenterology, 2010. 105(3): p. 493.

14. Leung, W. and J. Sung, Intestinal metaplasia and gastric carcinogenesis. Alimentary pharmacology & therapeutics, 2002. 16(7): p. 1209–1216.

15. Jencks, D.S., et al., Overview of Current Concepts in Gastric Intestinal Metaplasia and Gastric Cancer. Gastroenterol Hepatol (N Y), 2018. 14(2): p. 92–101.

16. Saw, P.E., J. Chen, and E. Song, Targeting CAFs to overcome anticancer therapeutic resistance. Trends Cancer, 2022. 8(7): p. 527–555.

17. Asif, P.J., et al., The role of cancer-associated fibroblasts in cancer invasion and metastasis. Cancers, 2021. 13(18): p. 4720.

18. Feng, B., et al., Cancer-associated fibroblasts and resistance to anticancer therapies: Status, mechanisms, and countermeasures. Cancer Cell International, 2022. 22(1): p. 166.

19. Ohlund, D., et al., Distinct populations of inflammatory fibroblasts and myofibroblasts in pancreatic cancer. J Exp Med, 2017. 214(3): p. 579–596.

20. Elyada, E., et al., Cross-Species Single-Cell Analysis of Pancreatic Ductal Adenocarcinoma Reveals Antigen-Presenting Cancer-Associated Fibroblasts. Cancer Discov, 2019. 9(8): p. 1102–1123.

21. Hu, B., et al., Subpopulations of cancer-associated fibroblasts link the prognosis and metabolic features of pancreatic ductal adenocarcinoma. Ann Transl Med, 2022. 10(5): p. 262.

22. Sahai, E., et al., A framework for advancing our understanding of cancer-associated fibroblasts. Nature Reviews Cancer, 2020. 20(3): p. 174–186.

23. Geng, X., et al., Cancer-Associated Fibroblast (CAF) Heterogeneity and Targeting Therapy of CAFs in Pancreatic Cancer. Front Cell Dev Biol, 2021. 9: p. 655152.

24. Jass, J.R., Role of intestinal metaplasia in the histogenesis of gastric carcinoma. J Clin Pathol, 1980. 33(9): p. 801–10.

25. Park, Y.H. and N. Kim, Review of atrophic gastritis and intestinal metaplasia as a premalignant lesion of gastric cancer. Journal of cancer prevention, 2015. 20(1): p. 25.

26. Tatematsu, M., T. Tsukamoto, and K. Inada, Stem cells and gastric cancer: role of gastric and intestinal mixed intestinal metaplasia. Cancer science, 2003. 94(2): p. 135–141.

27. Gipson, I.K., Goblet cells of the conjunctiva: A review of recent findings. Prog Retin Eye Res, 2016. 54: p. 49–63.

28. Van Landeghem, L., et al., Activation of two distinct Sox9-EGFP-expressing intestinal stem cell populations during crypt regeneration after irradiation. Am J Physiol Gastrointest Liver Physiol, 2012. 302(10): p. G1111–32.

29. Koulis, A., et al., CD10 and Das1: a biomarker study using immunohistochemistry to subtype gastric intestinal metaplasia. BMC Gastroenterol, 2022. 22(1): p. 197.

30. Hopkins, E.G., et al., Intestinal epithelial cells and the microbiome undergo swift reprogramming at the inception of colonic Citrobacter rodentium infection. MBio, 2019. 10(2): p. e00062–19.

31. Wang, J., et al., Differential gene expression in normal esophagus and Barrett’s esophagus. Journal of gastroenterology, 2009. 44: p. 897–911.

32. Takan, I., et al., “In the light of evolution:” keratins as exceptional tumor biomarkers. PeerJ, 2023. 11: p. e15099.

33. Aguilar-Medina, M., et al., SOX9 stem-cell factor: clinical and functional relevance in cancer. Journal of oncology, 2019. 2019.

34. Cao, W., et al., Claudin18. 2 is a novel molecular biomarker for tumor-targeted immunotherapy. Biomarker Research, 2022. 10(1): p. 1–21.

35. Lv, J. and P. Li, Mesothelin as a biomarker for targeted therapy. Biomarker Research, 2019. 7(1): p. 1–18.

36. Chu, Y., et al., Pan-cancer T cell atlas links a cellular stress response state to immunotherapy resistance. Nat Med, 2023. 29(6): p. 1550–1562.

37. Ostroumov, D., et al., CD4 and CD8 T lymphocyte interplay in controlling tumor growth. Cellular and molecular life sciences, 2018. 75: p. 689–713.

38. Echarti, A., et al., CD8+ and regulatory T cells differentiate tumor immune phenotypes and predict survival in locally advanced head and neck cancer. Cancers, 2019. 11(9): p. 1398.

39. Hao, D., et al., The Single-Cell Immunogenomic Landscape of B and Plasma Cells in Early-Stage Lung Adenocarcinoma. Cancer Discov, 2022. 12(11): p. 2626–2645.

40. Xue, R., et al., Liver tumour immune microenvironment subtypes and neutrophil heterogeneity. Nature, 2022. 612(7938): p. 141–147.

41. Cheng, S., et al., A pan-cancer single-cell transcriptional atlas of tumor infiltrating myeloid cells. Cell, 2021. 184(3): p. 792–809. e23.

42. Glabman, R.A., P.L. Choyke, and N. Sato, Cancer-associated fibroblasts: Tumorigenicity and targeting for cancer therapy. Cancers, 2022. 14(16): p. 3906.

43. Gascard, P. and T.D. Tlsty, Carcinoma-associated fibroblasts: orchestrating the composition of malignancy. Genes & development, 2016. 30(9): p. 1002–1019.

44. Mhaidly, R. and F. Mechta-Grigoriou. Fibroblast heterogeneity in tumor micro-environment: Role in immunosuppression and new therapies. in Seminars in immunology. 2020. Elsevier.

45. LeBleu, V.S. and R. Kalluri, A peek into cancer-associated fibroblasts: origins, functions and translational impact. Disease models & mechanisms, 2018. 11(4): p. dmm029447.

46. Mao, X., et al., Crosstalk between cancer-associated fibroblasts and immune cells in the tumor microenvironment: new findings and future perspectives. Molecular cancer, 2021. 20(1): p. 1–30.

47. Liu, T., et al., Cancer-associated fibroblasts: an emerging target of anti-cancer immunotherapy. Journal of hematology & oncology, 2019. 12(1): p. 1–15.

48. Sinjab, A., et al., Resolving the Spatial and Cellular Architecture of Lung Adenocarcinoma by Multiregion Single-Cell Sequencing. Cancer Discov, 2021. 11(10): p. 2506–2523.

49. Galletti, G., et al., Two subsets of stem-like CD8+ memory T cell progenitors with distinct fate commitments in humans. Nature immunology, 2020. 21(12): p. 1552–1562.

