## Supplementary Information for "METI: Deep profiling of tumor ecosystems by integrating cell morphology and spatial transcriptomics"

**Supplementary Table S1. Datasets analyzed in this paper**

| Species | Tissue | Dataset dimensions | Protocol | Spot diameter by pixels |
| --- | --- | --- | --- | --- |
| Human | Stomach Adenocarcinoma-G1 | 1202 spots  17943 genes | 10x Visium | 72 pixels |
| Human | Stomach Adenocarcinoma-G2 | 4328 spots  20615 genes | 10x Visium | 59 pixels |
| Human | Stomach Adenocarcinoma-G3 | 3875 spots  17943 genes | 10x Visium | 130 pixels |
| Human | Stomach Adenocarcinoma-G4 | 4130 spots  20615 genes | 10x Visium | 59 pixels |
| Human | Lung adenocarcinoma-L1 | 3360 spots 17943 genes | 10x Visium | 258 pixels |
| Human | Bladder Cancer-B1 | 9029 spots  18085 genes | 10x Visium | 236 pixels |
| Human | Bladder Cancer-B2 | 9963 spots 18085 genes | 10x Visium | 238 pixels |

**Supplementary Table S2. Specific marker genes used for cell type detection.**

| Cell type | Marker genes |
| --- | --- |
| Goblet cells | *MS4A10, MGAM, CYP4F2, XPNPEP2, SLC5A9, SLC13A2, SLC28A1, MEP1A, ABCG2, ACE2* |
| Tumor cells | *EPCAM, SOX9, CLDN18, MSLN* |
| T cells | *CD3D, CD3E* |
| CD4 T cells | *CD4* |
| CD8 T cells | *CD8A, CD8B* |
| Tregs | *FOXP3,* *IL2RA* |
| Tex | *HAVCR2, LAG3, CTLA4, TIGIT, PDCD1*, *LAYN* |
| Neutrophils | *HMGB2*, *IFIT1*, *ISG15*, *LCN2*, *MX1*, *S100A8*, *S100A9*, *SYAP1* |
| B cells | *CD19, MS4A1* |
| Plasma cells | *MZB1, JCHAIN* |
| Macrophages | *CD4*, *CD80*, *CD86*, *CCR5*, *CD14*, *CD163*, *LILRB4*, *CD33*, *TLR2*, *TLR4* |
| Fibroblasts | *COL1A1, COL1A2, COL6A1, COL6A2* |
| myCAFs | *TGFB1, ACTA2* |
| iCAFs | *CXCL12, CXCL13, CXCL14* |
| apCAFs | CD74, *HLA-DQA1, HLA-DPB1* |

**Supplementary Table S3. Software mentioned in this paper**.

| **Method** | **Version** | **URL** | **Reference** |
| --- | --- | --- | --- |
| TESLA | 1.2.4 | https://github.com/jianhuupenn/TESLA | Hu *et al*.[1] |
| SpaGCN | 1.2.7 | https://github.com/jianhuupenn/SpaGCN | Hu *et al*.[2] |
| RCTD | 1.2.0 | https://github.com/dmcable/RCTD | Cable *et al*.[3] |
| MUSE | 0.0.8 | https://github.com/AltschulerWu-Lab/MUSE | Bao *et al*.[4] |
| CellTrek | 0.0.94 | https://github.com/navinlabcode/CellTrek | Wei *et al*.[5] |
| CytoSPACE | 1.0.6 | https://github.com/digitalcytometry/cytospace | Vahid *et al*.[6] |

**Supplementary Figure S1.** Overlay of regions expressing tumor-related genes and tumor specific markers positive regions, including MKI67, MSLN, and CLDN18.

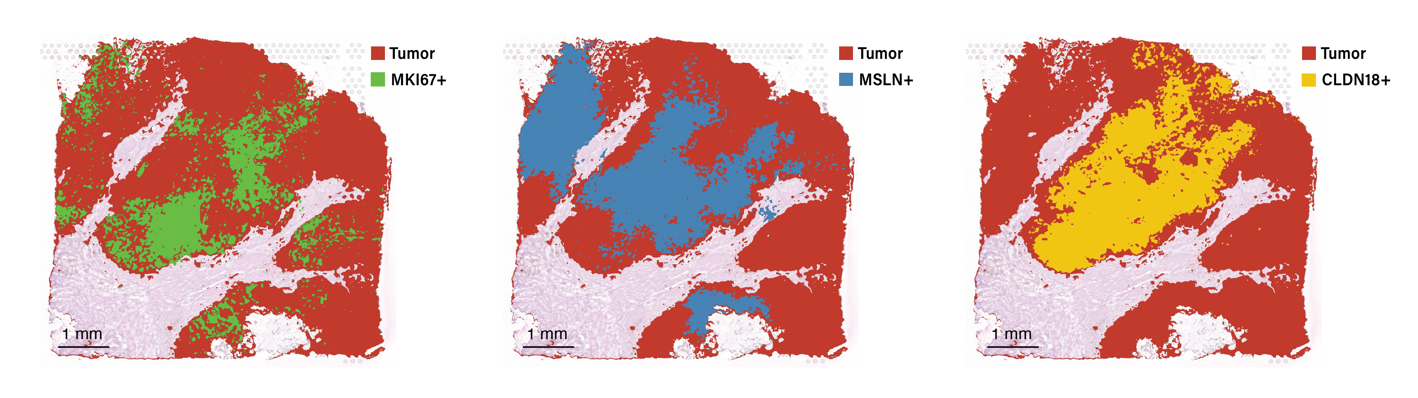

**Supplementary Figure S2.** Pathology annotation of Lymphoid aggregate enriched spots on STAD-G4 sample and macrophages enriched spots on LUAD-L1 sample.

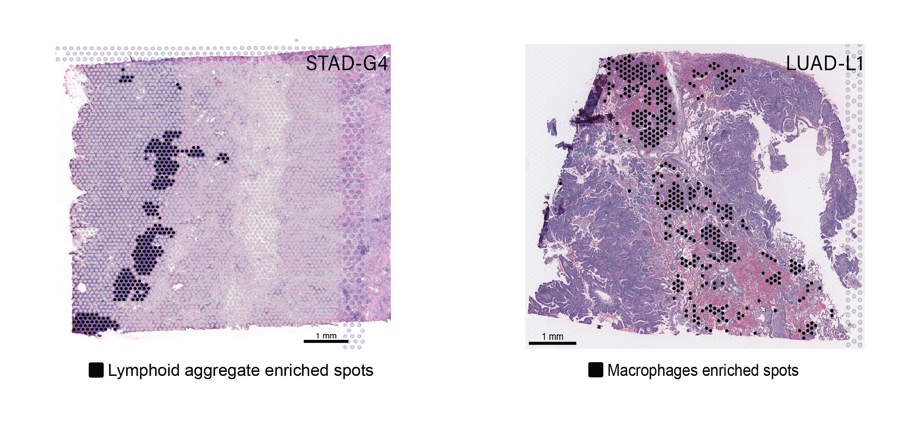

b

a

**Supplementary Figure S3.** Pathology annotation of neutrophils enriched spots on BLCA-B1 and BLCA-B2 samples.

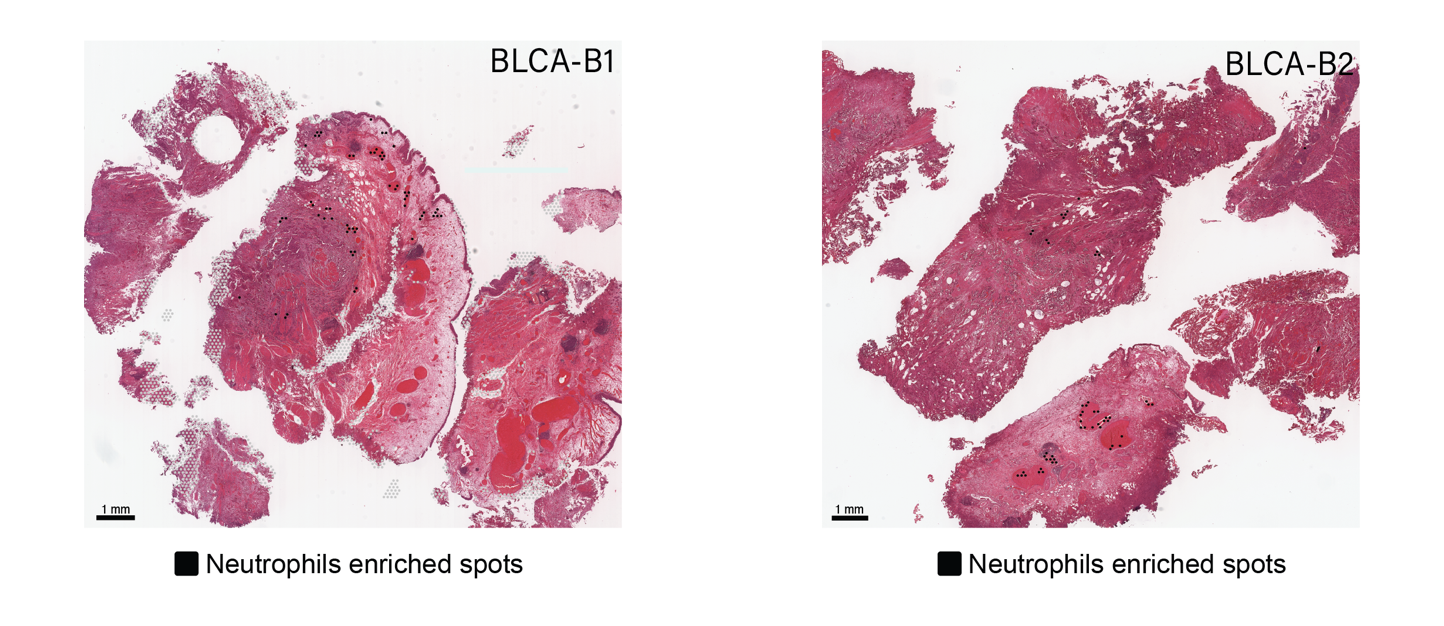

**References**

1.	Hu, J., et al., *Deciphering tumor ecosystems at super resolution from spatial transcriptomics with TESLA.* Cell Systems, 2023. **14**(5): p. 404-+.

2.	Hu, J., et al., *SpaGCN: Integrating gene expression, spatial location and histology to identify spatial domains and spatially variable genes by graph convolutional network.* Nat Methods, 2021. **18**(11): p. 1342-1351.

3.	Cable, D.M., et al., *Robust decomposition of cell type mixtures in spatial transcriptomics.* Nat Biotechnol, 2022. **40**(4): p. 517-526.

4.	Bao, F., et al., *Integrative spatial analysis of cell morphologies and transcriptional states with MUSE.* Nat Biotechnol, 2022. **40**(8): p. 1200-1209.

5.	Wei, R., et al., *Spatial charting of single-cell transcriptomes in tissues.* Nat Biotechnol, 2022. **40**(8): p. 1190-1199.

6.	Vahid, M.R., et al., *High-resolution alignment of single-cell and spatial transcriptomes with CytoSPACE.* Nat Biotechnol, 2023.
